# From asexuality to sexual reproduction: cyclical switch of gametogenic pathways in hybrids depends on ploidy level

**DOI:** 10.1101/2023.06.18.545483

**Authors:** Dmitrij Dedukh, Anatolie Marta, Ra-Yeon Myung, Myeong-Hun Ko, Da-Song Choi, Yong-Jin Won, Karel Janko

## Abstract

The cellular and molecular mechanisms governing sexual reproduction is highly conserved across eukaryotes. Nevertheless, hybridization can disrupt such machinery leading to asexual reproduction. To investigate how hybridization and polyploidization affect gametogenesis and reproductive outcomes of asexual hybrids, we conducted a comprehensive study on diploid and triploid hybrids along with their sexual parental species from the freshwater fish family Cobitidae. In diploid and triploid hybrids, most gonocytes maintain their original ploidy level. During meiosis, such gonocytes experience abnormal chromosome pairing preventing progression beyond pachytene. Diploid hybrid females regain fertility through premeiotic genome endoreplication, resulting in the rare emergence of tetraploid gonocytes. Tetraploid gonocytes bypass meiosis and lead to clonal diploid gametes. In contrast, triploid hybrids lack genome endoreplication but utilize premeiotic genome elimination of a single-copy parental genome forming diploid gonocytes that undergo meiosis and produce haploid gametes. Therefore, the interplay of parental genomes leads to diverse gametogenic outcomes in hybrids dependent on their ploidy and genome dosage. These alterations in gametogenic pathways can persist across generations, potentially enabling the cyclic maintenance of asexual/polyploid hybrids in natural populations.

## Introduction

Sexual reproduction is the almost universal feature of eukaryotes and includes the meiotic formation of gametes with a half genome of diploid, fertilization and development of new organisms (Lenormand et al., 2016; Otto and Lenormand, 2002). The cellular and molecular machinery ensuring gametogenesis, recombination and fertilization is primarily conserved (Lenormand et al., 2016). However, interspecific hybridization may disrupt the usual reproductive pathways, reducing fertility in hybrids (Arnold and Hodges, 1995; Coyne et al., 2004; Rieseberg, 2001). Nevertheless, hybridization may also lead to the emergence of various types of so-called asexual reproduction among different plant and animal species (Abbott et al., 2013; Dawley and Bogart, 1989; Janko et al., 2018; Schön et al., 2009; Stöck et al., 2021). Forms of asexual reproduction have traditionally been categorized based on types of produced gametes and whether sperm is required for their development (Dawley and Bogart, 1989; Schön et al., 2009; Stöck et al., 2021). Parthenogenesis refers to the production of unreduced gametes which spontaneously develop into usually clonal progeny (Dawley and Bogart, 1989; Schön et al., 2009; Stöck et al., 2021). Gynogenesis (or sperm-dependent parthenogenesis) involves the formation of clonal gametes which require sperm to trigger their development without karyogamy (Dawley and Bogart, 1989; Dedukh and Krasikova, 2022; Schön et al., 2009; Stöck et al., 2021). Hemi- or mero-clonal reproduction involve kleptogenesis and hybridogenesis, where part of asexual’s genome is eliminated, while the other part is passed to gametes which requires fertilization with karyigamy (Bogart et al., 2007; Dawley and Bogart, 1989; Dedukh and Krasikova, 2022; Schön et al., 2009; Stöck et al., 2021). Interestingly, experiments have demonstrated that shifts to asexuality occur directly in the F1 generation of some hybrids meaning that asexual gametogenesis do not require novel pathways to evolve, but may rely on misregulation of cellular mechanisms present in sexual parental species (Albertini et al., 2019; Brownfield and Köhler, 2011; Carman, 1997; Choleva et al., 2012; Dedukh et al., 2019a; Marta et al., 2023; Mason and Pires, 2015).

At cellular level, the situation is even more complex as fundamentally different cellular and molecular mechanisms may generate the same output, such as the production of clonal gamete (Dedukh and Krasikova, 2022; Stöck et al., 2021). For instance, clonal gametogenesis may be achieved by premeiotic endoreplication that causes ploidy elevation during gonocyte proliferation e.g., in diploid hybrids, the chromosome set of gonocytes becomes tetraploid (Cimino, 1972b; Dedukh et al., 2022a; Dedukh et al., 2022b; Dedukh et al., 2020b; Itono et al., 2006; Kuroda et al., 2018; Lutes et al., 2010; Macgregor and Uzzell, 1964; Stöck et al., 2002). Such tetraploid gonocytes proceed to supposedly normal meiosis, but pairing occurs only between duplicated copies of the same chromosomes, thereby delivering no variability among progeny (Dedukh et al., 2022a; Dedukh et al., 2020b; Janko et al., 2021; Kuroda et al., 2018; Lutes et al., 2010; Macgregor and Uzzell, 1964). A contrasting mechanism involves achiasmatic meiosis, as in hybrid *Poecilia formosa*, where the ploidy level of gonocytes remains unmodified but chromosomes exist as univalents during the first meiotic prophase with no sign of recombination (Dedukh et al., 2022b; Monaco et al., 1984). Hypothetically, the reductional division is skipped, but the equational division is normal again resulting in the same type of clonal progeny (Dedukh et al., 2022b; Monaco et al., 1984).

After forming clonal eggs, asexuals have to avoid fertilization to prevent ploidy elevation, as polyploid animals are usually sterile. In gynogenetic organisms, sperm is required to activate the egg but not incorporate its genetic material into the egg (Beukeboom and Vrijenhoek, 1998; Zhang et al., 2015). To maintain clonal lineages in natural populations, gynogens thus have to exploit males of sexual species, whose sperm is “wasted”, acting as so-called “sexual parasites” (Alves et al., 2001; Bi and Bogart, 2010; Gu et al., 2022; Majtánová et al., 2016; Morishima et al., 2008a). Nevertheless, occasional failure to eliminate sperm pronucleus from the zygote leads to the emergence of triploid offspring (Alves et al., 1998; Lamatsch and Stöck, 2009; Zhang et al., 2015). Triploids were generally assumed to maintain the same type of gynogenetic reproductive mode as their diploid hybrid progenitors (Bogart et al., 2007; Cuellar, 1971; Dedukh et al., 2021; Dedukh et al., 2020b; Lutes et al., 2010; Monaco et al., 1984), but some can switch reproductive modes and exploit genome elimination during gametogenesis (Alves et al., 1998; Cimino, 1972a; Cimino, 1972b; Goddard et al., 1998; Kim and Lee, 1990; Saitoh et al., 2004). For instance, during the so-called triploid or meiotic hybridogenesis, a single-copied genome is eliminated (for example, AAB hybrids between parental species A and B eliminate the genome B), while the double-copied genomes (AA in AAB hybrids) enters normal meiosis and undergoes pairing and recombination, resulting in haploid gametes (Alves et al., 1998; Christiansen and Reyer, 2009; Dedukh et al., 2015; Goddard et al., 1998; Kim and Lee, 1990; Saitoh et al., 2004; Stöck et al., 2012). The examples above show that the transition between sexual and asexual modes (and back) may be pretty dynamic. It also frequently depends on the ploidy level of the individuals, when the combination of genomes from the same parental species may deliver very different gametogenic output in diploid and triploid hybrids.

However, underlying cytogenetic mechanisms are unknown for most of asexual hybrid complexes (Alves et al., 1998; Christiansen and Reyer, 2009; Dedukh et al., 2015; Goddard et al., 1998; Kim and Lee, 1990; Saitoh et al., 2004; Stöck et al., 2012). Additionally, cellular and molecular machinery causing such alterations between reproduction types remains even less understood. Detailed investigation of gametogenic pathways in asexuals should therefore answer basic questions like what mechanisms cause the transitions from sexual reproduction to asexuality? Why some types of gametogenic aberrations are more common than others? And do similar gametogenic alterations rely on similar cellular and molecular mechanisms in different asexuals?

A suitable model to address such questions in detail is the *Cobitis hankugensis-Iksookimia longicorpa* hybrid complex of Korean loaches (Kim and Lee, 1990; Lee, 1992; Lee, 1995; Saitoh et al., 2004; Ko, 2009). In this complex (formerly reported as *C. sinensis-longicorpus* or *C. hankugensis-Iksookimia longicorpus*), two diploid parapatrically distributed bisexual species, *C. hankugensis* (HH, 2n = 48 chromosomes) and *I. longicorpa* (LL, 2n = 50 chromosomes) meet in a hybrid zone and form diploid hybrids (HL, 2n = 49 chromosomes) (Figure 1) (Kim and Lee, 1990; Kim et al., 2000; Saitoh et al., 2004; Ko, 2009). From previous reproductive experiments it was hypothesized that diploid hybrids produced diploid clonal gametes, which incorporate the sperm genome of one of the parental species and form triploid organisms with two genome compositions, namely HHL (3n=73 chromosomes) and LLH (3n=74 chromosomes) (Figure 1) (Lee, 1995; Saitoh et al., 2004; Ko, 2009). Triploid females, by contrast, produce either HL diploid hybrid progeny, of diploid HH and LL progeny, depending on which parental species fertilized their eggs (Figure 1) (Kim and Lee, 1990; Lee, 1995; Saitoh et al., 2004; Ko, 2009). This indicates that triploids may possess genome elimination during their gametogenesis and probably form recombined haploid gametes. Nevertheless, gametogenic mechanisms underlying the formation of unreduced gametes in diploid hybrids and putatively haploid gametes in triploid hybrids remain unknown.

**Figure 1.**
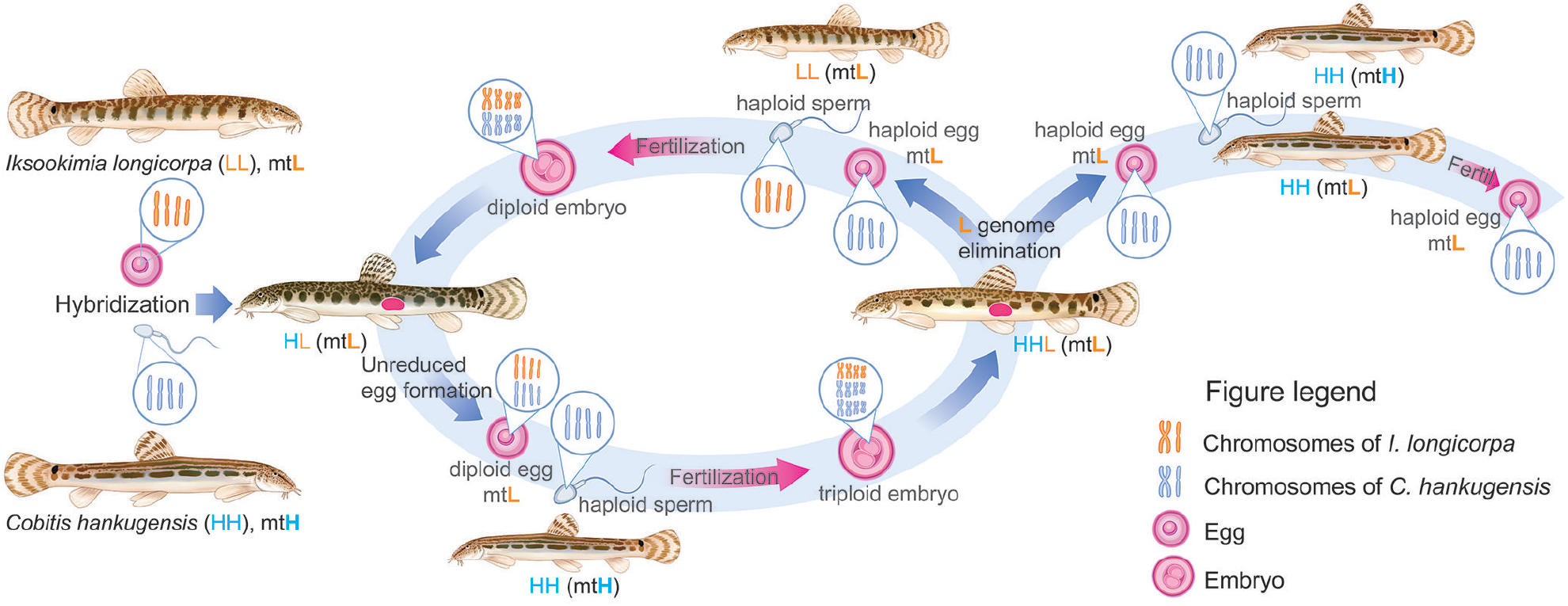
Schematic overview of gametogenesis and reproduction of diploid and triploid hybrids within *C. hankugensis-I. longicorpa* complex. (Redrawn from (Kim et al., 2000; Saitoh et al., 2004; Ko, 2009). After crosses of two parental sexual species, *Cobitis hankugensis* (HH, marked in blue) and *Iksookimia longicorpa* (LL, marked in orange), diploid hybrids (HL) are produced with the mitochondrial DNA (designated as ‘mt’) from one of the sexual species (L). Diploid hybrids form diploid clonal gametes with ‘L’ mtDNA. After fertilization of such eggs by sperm of one of the parental species, triploid hybrids with L mtDNA appear (HHL). In triploids, a single-copied genome (L) is eliminated during their gametogenesis, and the remaining haploid gametes (HH) produce haploid ‘H’ gametes with ‘L’ mtDNA. After fertilization of such gametes by sperm from *C. hankugensis*, diploid sexual species appear but with ‘L’ mtDNA. After fertilization of gametes produced by triploids by sperm from the other parental species, *I. longicorpa*, new diploid hybrids appear with ‘L’ mtDNA.

Our study investigates particular cellular mechanisms in natural diploid and triploid hybrids with different reproductive outcomes. We examined oocytes during the pachytene and diplotene stages of meiosis and the gonocyte’s genome composition and ploidy level. Additionally, we analysed the distribution of meiocytes throughout the ovaria of asexual diploid and triploid hybrid biotypes. Finally, we investigated the mechanisms of hybrid sterility in triploid hybrid males.

## Results

### Sexual species exhibit normal pairing of chromosomes

We found that the somatic cells of *C. hankugensis* and *I. longicorpa* have 2n = 48 and 2n = 50 chromosomes, respectively, identical to the previous finding (Kim and Lee, 1990). Thus, in the somatic cells of diploid HL hybrids, we observed 2n = 49 chromosomes; in the somatic cells of triploid HHL hybrids, we detected 3n = 73, which was also the same as the previous finding (Kim and Lee, 1990). FISH with distinguished satellite DNA SatCE1 and SatCE5 repeat markers (Marta et al., 2020) showed the presence of staining in two small submetacentric chromosomes in both sexual species (Supplementary Figure S1A, B) and in three chromosomes of triploid HHL hybrids (Supplementary Figure S1C). Thus, FISH based mapping of chromosome specific SatCE1 tandem repeat marker may serve as a reliable tool to identify parental genomes in hybrid individuals (Supplementary Figure S1C).

We investigated pachytene chromosomes in three males and one female of *C. hankugensis* and one male and three females of *I. longicorpa* (Supplementary Table 1). To confirm bivalent formation during the pachytene stage of the sexual species, we stained the axial (SYCP1) and lateral (SYCP3) elements of the synaptonemal complexes. In males and females of both parental species, we observed the same number of chromosomes as in their somatic cells, paired into bivalents with no univalent or aberrant pairing. In males and females of *C. hankugensis*, we detected 24 bivalents, and in males and females of *I. longicorpa*, we observed 25 bivalents (Figure 2A, Supplementary Figure S2A-C). On pachytene spreads of both males and females, we visualized crossing over loci and detected at least one signal per each bivalent. In males of *C. hankugensis* and *I. longicorpa*, we usually observed distal localization of MLH1 loci in contrast to females, where we usually detected interstitial localization (Figure 3A, Supplementary Figure S3A-B). To confirm the results of pachytene analysis, we analyzed diplotene oocytes and found 24 bivalents in *C. hankugensis* females and 25 bivalents in *I. longicorpa* females (Supplementary Figure S4A, B). Bivalents in diplotene oocytes are united with chiasmata, which correspond to crossover loci. To confirm the pairing between homologous chromosomes in both parental species, we performed FISH with SatCE1 marker. We observed a signal on each chromosome in a particular bivalent in diplotene chromosomal spreads of *C. hankugensis* and *I. longicorpa* (Supplementary Figure S5A, B).

**Figure 2.**
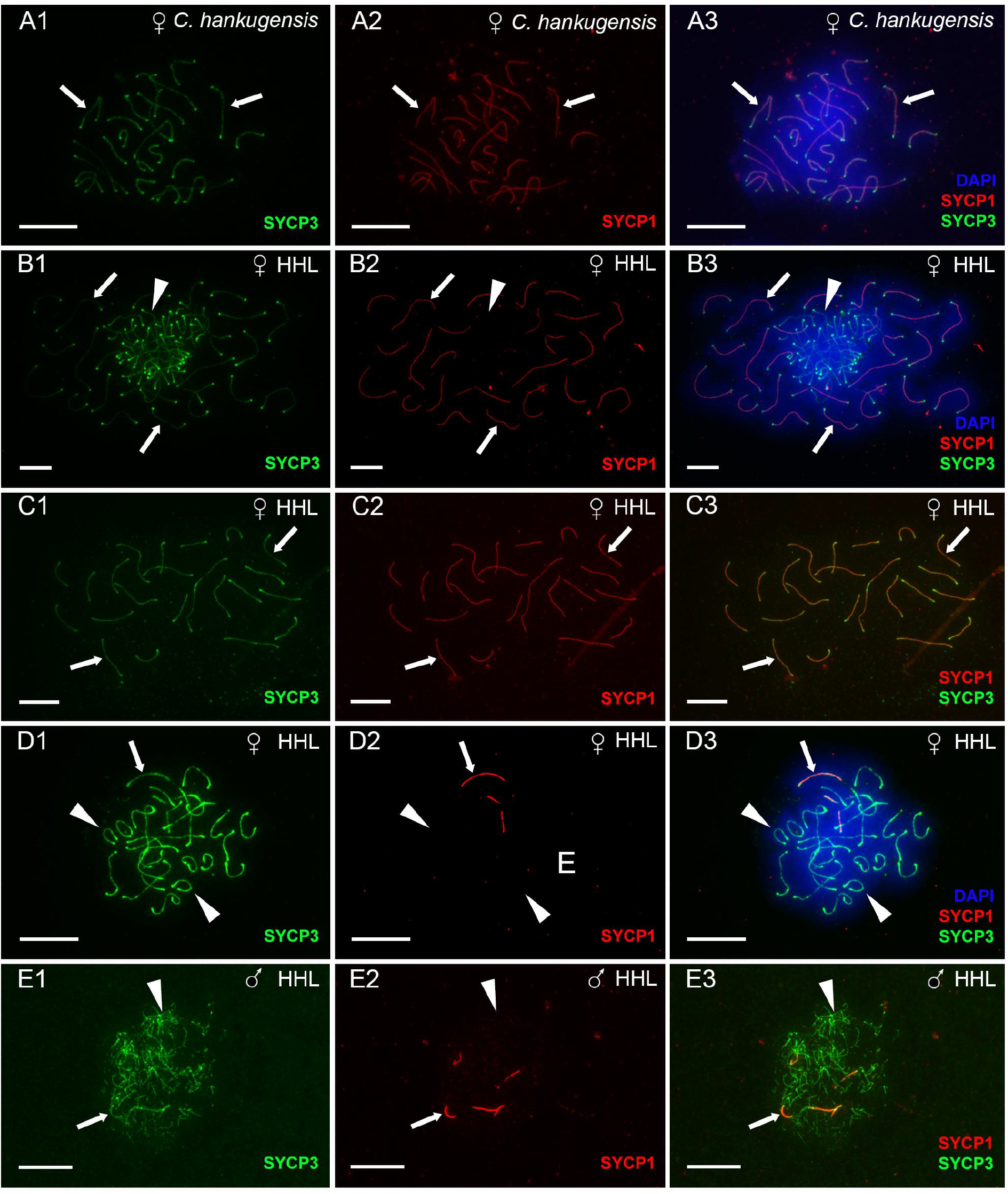
The analysis of pairing in pachytene oocytes (A1-D3) and spermatocytes (E1-E3) of *C. hankugensis* (A1-A3) and triploid HHL hybrids (B1-E3). Synaptonemal complexes were visualized using immunostaining of lateral (SYCP3 protein, green) (A1, B1, C1, D1, and E1) and central (SYCP1 protein, red) (A2, B2, C2, D2, and E2) components. Corresponding merged figures (A3, B3, C3, D3, and E3) also include DAPI staining (blue). Accumulation of SYCP3 and SYCP1 proteins (indicated by thick arrows) allows distinguishing bivalents, while univalents accumulate only SYCP3 protein (indicated by arrowheads). Pachytene oocytes of *C. hankugensis* exhibit 24 fully paired bivalents (A1–A3). In triploid hybrids, we observed pachytene oocytes with 24 bivalents and 25 univalents (B1–B3), oocytes with 24 bivalents (C1– C3), and oocytes with 25 univalents (D1–D3). Triploid hybrid males exhibit pachytene oocytes only with the aberrant pairing of several bivalents and univalent (E1–E3). Scale bar = 10 µm.

**Figure 3.**
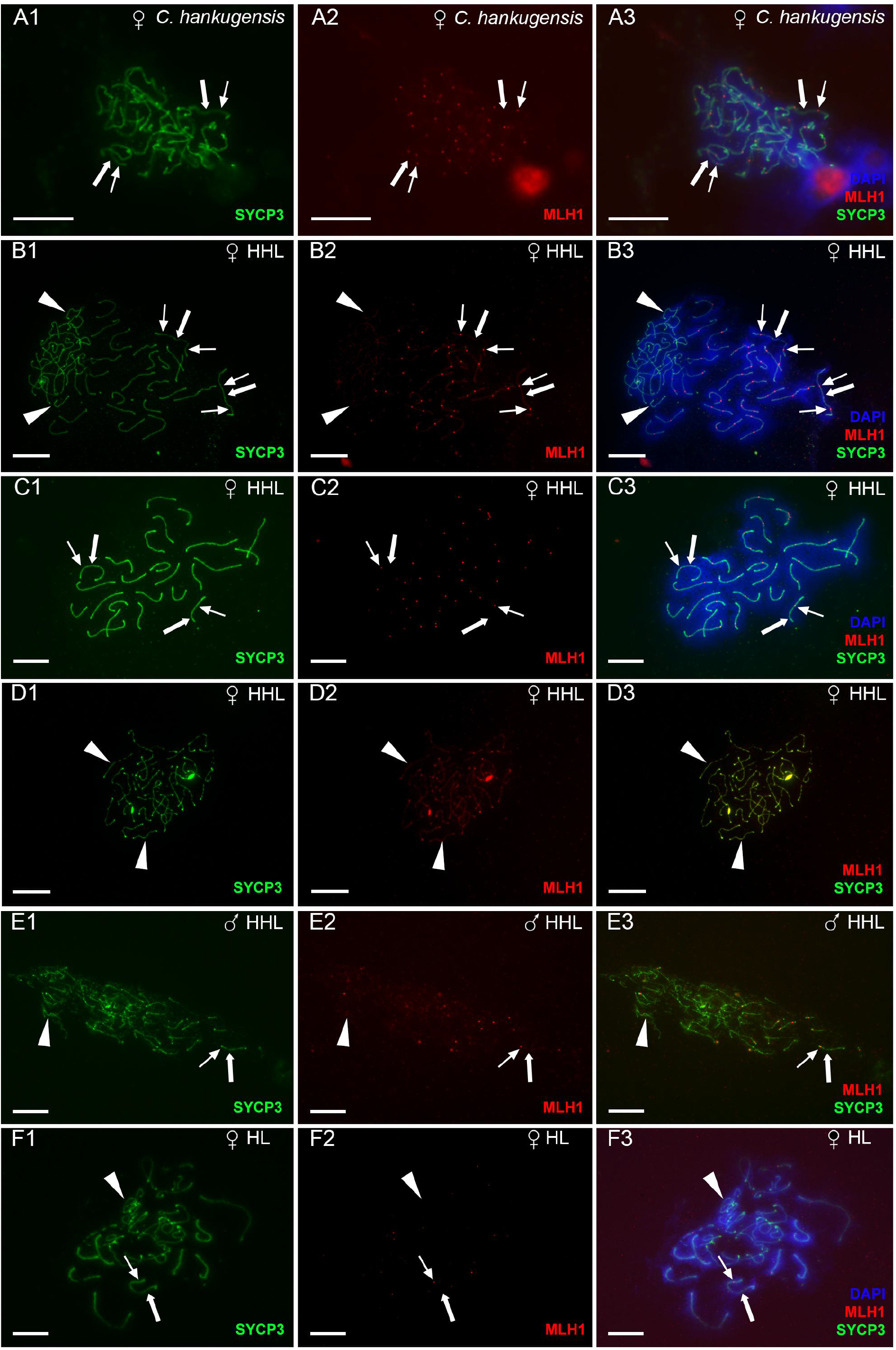
The analysis of crossover loci in pachytene oocytes (A1-D3, F1-F3) and spermatocytes (E1-E3) from gonads of *C. hankugensis* (A1-A3), triploid HHL hybrids (B1-E3) and diploid hybrid (F1-F3). Crossover loci were detected by MLH1 protein (indicated by thin arrows, red) (A2, B2, C2, D2, E2, and F2) on lateral components of synaptonemal complexes (SYCP3 protein, green) (A1, B1, C1, D1, E1, and F1). Corresponding merged figures (A3, B3, C3, D3, E3, and F3) also include DAPI staining (blue). MLH1 bindings (indicated by thin arrows, red) are located on bivalents (indicated by thick arrows) and do not accumulate on univalents (indicated by arrowheads). Pachytene oocytes of *C. hankugensis* exhibit 24 fully with at least one crossover locus per bivalent (A1–A3). In triploid hybrids, oocytes with 24 bivalents and 25 univalents have MLH1 signals only on bivalents (B1–B3). Oocytes with exclusively 24 bivalents (C1–C3) have recombination signals on each bivalent while oocytes with exclusively 25 univalents (D1–D3) do not have crossover locus. MLH1 immunostaining demonstrates the presence of crossover in individual bivalents formed in a triploid hybrid male (E1–E3) and pachytene oocytes with a unduplicated genome (F1–F3). Scale bar = 10 µm.

In addition, we investigated gonadal microanatomy and revealed the distribution of gonocytes, meiocytes and gametes in both males and females of the studied sexual species (Supplementary Figure S6A-D). In *C. hankugensis* and *I. longicorpa* females, we observed gonocytes and pachytene clusters between pre-vitellogenic and vitellogenic oocytes. In *C. hankugensis* and *I. longicorpa* males, we detected different clusters of gonocytes, spermatocytes during pachytene and spermatocytes during metaphase I, and large clusters of spermatids. The morphology of their nuclei discriminated different cell types after DAPI staining according to the previously published results for *Cobitis* species (Dedukh et al., 2021; Dedukh et al., 2020b; Marta et al., 2023).

We also identified the ploidy of gonocytes, pachytene oocytes and early diplotene oocytes in sexual species using whole mount FISH with satDNA marker SatCE1 (Supplementary Figure S7). In gonocytes (n = 23) of sexual species, we distinguished two signals suggesting their diploid genome composition (Supplementary Figure S7C). We observed one large signal in pachytene cells (n = 98) as chromosomes formed bivalents (Supplementary Figure S7B). In small (nucleus diameter 8-15 mkm; n = 68) and larger (nucleus diameter 15-40 mkm; n = 93) diplotene oocytes, we distinguished two adjacent signals indicating the chromosome separation but still connected by chiasmata, which are clearly visible on later stages during lampbrush chromosome analysis (Supplementary Figure S7A).

### Diploid hybrid females exploit premeiotic genome duplication and produce unreduced eggs

To identify a gametogenic pathway in diploid hybrids, we first determined the number of bivalents in their pachytene and diplotene oocytes (Supplementary Table 1). After analysis of 36 diplotene oocytes from one diploid hybrid female, we detected only oocytes with 49 bivalents, suggesting that the number of chromosomes in all analyzed oocytes was tetraploid (4n = 98 chromosomes) (Figure 4B). These results indicate the presence of premeiotic genome endoreplication during the gametogenesis of diploid hybrid females. FISH with SatCE1 DNA marker showed signals in both bivalents corresponding to *C. hankugensis* and *I. longicorpa* (Supplementary Figure S5D, E) suggesting the pairing between chromosomal copies emerged after premeiotic genome duplication.

**Figure 4.**
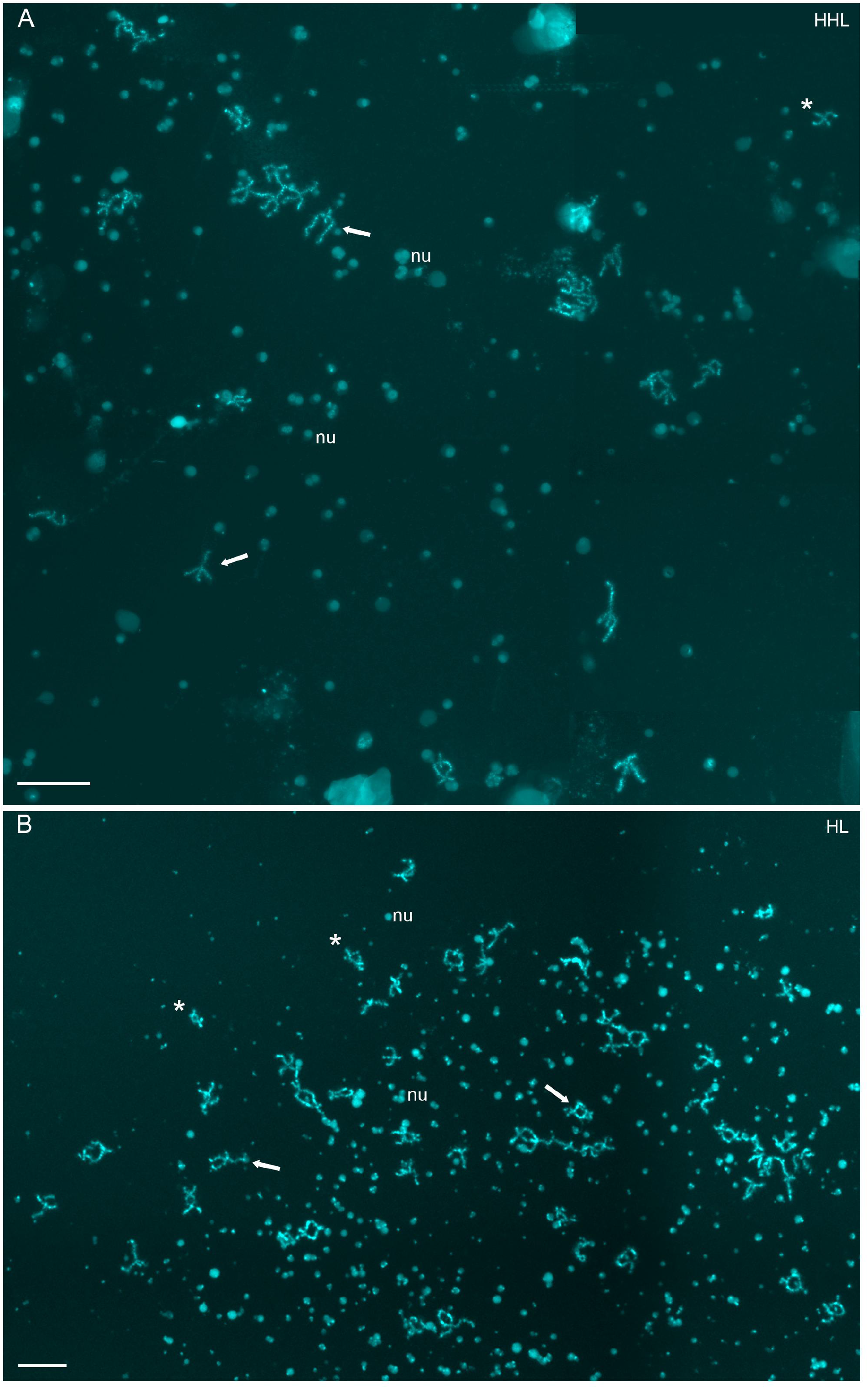
Diplotene chromosomal spreads from the individual oocytes of triploid HHL (A) and diploid HL (B) hybrid females. A triploid hybrid’s chromosomal set of diplotene oocytes includes 24 bivalents, possibly of *C. hankugensis* (A). The chromosomal set of diploid hybrid includes 49 bivalents (B). Since the chromosomal spread from the individual oocyte was large, four images were merged into one in the case of A and B. Chromosomes were stained with DAPI (cyan). Thick arrows indicate examples of individual bivalents; nu shows examples of extrachromosomal nucleoli (nu). Asterisks indicate enlarged bivalents in Supplementary Figure S5C (HHL) and S5D and S5E (HL) for triploid and diploid hybrids, respectively. Scale bar = 50 µm.

In contrast to diplotene oocytes, in pachytene, we observed oocytes only with unduplicated genomes (Figure 3F). In total, we found 13 oocytes during pachytene from two hybrid females. Pachytene cells with unduplicated genomes exhibit aberrant pairing with 3-5 bivalents while other chromosomes exist as univalents (Figure 3F). This suggests that such oocytes possessed unduplicated genomes with 24 chromosomes of *C. hankugensis* and 25 chromosomes of *I. longicorpa*.

During analysis of gonadal microanatomy of hybrids reveal the presence of all cell types similar to parental species (Supplementary Figure S6G). To test whether genome endoreplication occurs premeiotically, we applied FISH with SatCE1 DNA marker to identify ploidy level in gonocytes, pachytene and early diplotene oocytes (Figure 5E, F). During the analysis of gonocytes in diploid HL hybrids, we detected cells with two signals (n = 297) and with four signals (n = 18), suggesting the presence of diploid and tetraploid gonocytes populations correspondingly (Figure 5F). We assume that tetraploid gonocytes emerged after genome endoreplication, corroborating the analysis of pachytene spreads. However, in pachytene, we cannot find diploid and tetraploid cell populations as both cell populations have two signals (two signals from univalents in diploid oocytes and two signals from paired bivalents in tetraploid oocytes). Nevertheless, we clearly observed two pairs of signals in early diplotene oocytes, suggesting that only duplicated oocytes can proceed beyond pachytene (Figure 5E).

**Figure 5.**
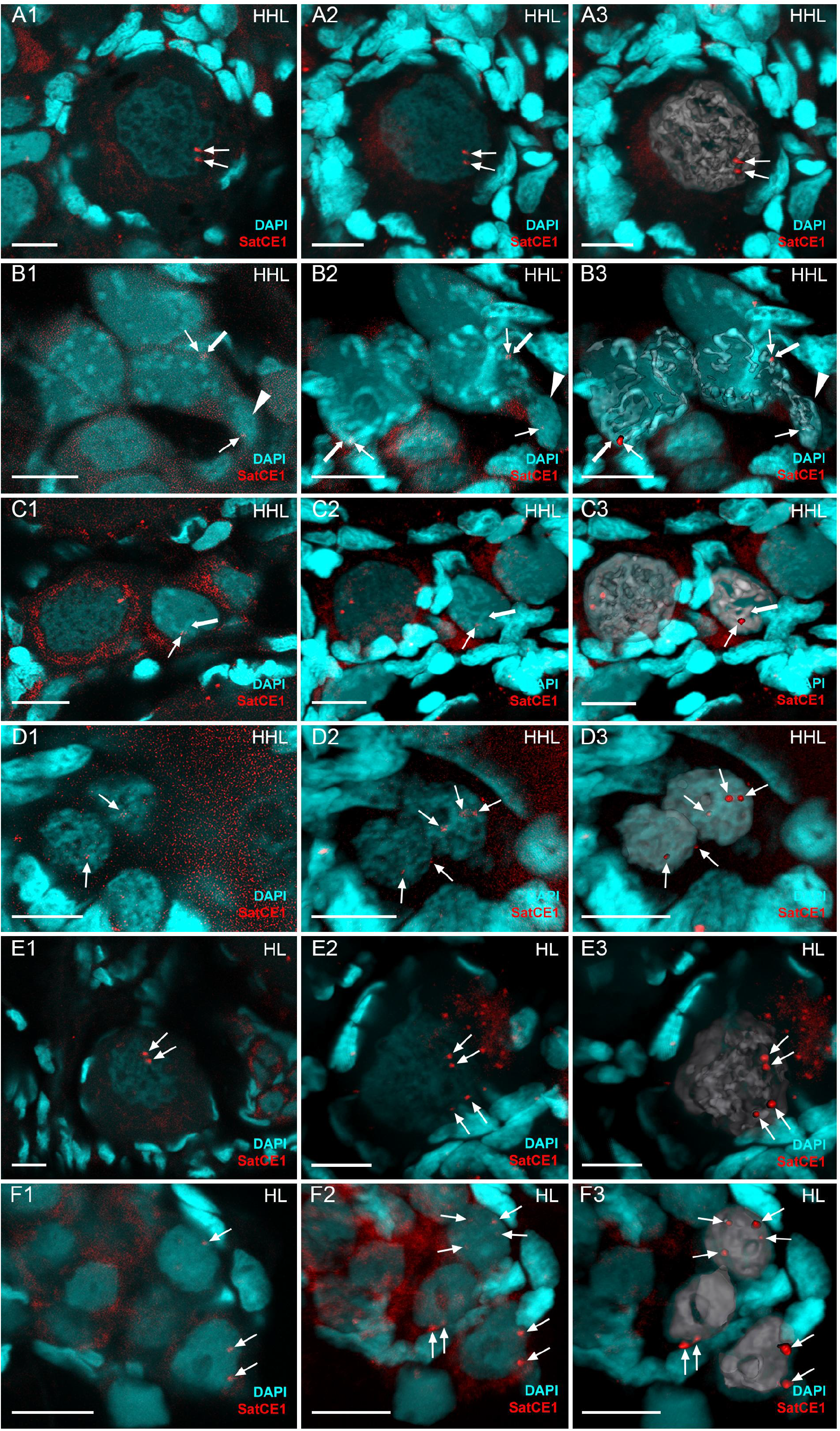
Identification of ploidy level of cells in gonadal fragments of triploid HHL (A1-D3) and diploid hybrids (E1-F3) using whole-mount FISH with chromosome-specific SatCE1 marker. In the diplotene oocyte of triploid HHL hybrid (A1–A3), two adjacent signals are visible suggesting the presence of two homologous chromosomes. Pachytene oocytes with bivalents and univalents (B1–B3) have signals on bivalent (indicated by thick arrow) as well as on univalent (indicated by arrowhead). Pachytene oocytes only with bivalents have one signal (indicated by arrow) on bivalent (indicated by thick arrow) (C1–C3). Diploid gonocytes with two signals (indicated by arrows) and triploid gonocytes with three signals (indicated by arrows) (D1–D3) in the ovary from triploid HHL hybrid. In the diplotene oocyte of diploid HL hybrid (E1–E3), two pairs of signals are visible, suggesting the presence of two bivalents. Diploid gonocyte with two signals (indicated by arrows) and tetraploid gonocyte with four signals (indicated by arrows) (F1–F3) in the ovary from diploid HL hybrid. DNA is stained by DAPI (cyan). Images (A1, B1, C1, D1, E1, and F1) are single confocal sections of 0.7 µm in thickness; corresponding 3D reconstructions (A2, B2, C2, D2, E2, and F2) and 3D surface reconstructions (A3, B3, C3, D3, E3, and F3) of metaphase plates with constructed isosurfaces of the signals and cells of interest. Scale bar = 10 µm.

We tested whether incidences of cells with endoreplicated genomes differ among diploid and triploid hybrids. To perform such analysis, we used the generalized linear model with binomial error distribution to compare the counts of gonocytes with initial ploidy level and duplicated ones. As a result, we found highly significant differences (p < 10^-4^), whereby diploids possesses ∼ 6% of duplicated cells, while HHL triploids had none.

### Triploid hybrid females perform premeiotic genome elimination and produce recombinant haploid eggs

Further, we investigated pachytene and diplotene oocytes to identify a gametogenic pathway in triploid hybrids (Supplementary Table S1). After analysis of 77 diplotene oocytes from seven triploid HHL hybrid females, we observed 24 bivalents possibly corresponding to the *C. hankugensis* genome (Figure 4A). We did not find univalent or abnormal pairing. These results suggest that *I. longicorpa* chromosomes were eliminated before the diplotene stage of meiosis, while two sets of *C. hankugensis* chromosomes form 24 bivalents. The presence of chiasmata between paired chromosomes confirms the incidence of recombination between putatively homologous chromosomes. FISH with SatCE1 DNA marker showed signals in two chromosomes from one bivalent, suggesting the pairing of homologous chromosomes (Supplementary Figure S5C).

By contrast, the analysis pachytene oocytes in six triploid HHL hybrid females revealed the presence of three cells populations differed in ploidy level (Supplementary Table S1). First population of cells (n = 153) included pachytene oocytes with initial ploidy level. In such oocytes we detected 24 bivalents possibly formed by *C. hankugensis* chromosome and 25 univalents perhaps representing *I. longicorpa* chromosomes clustered together (Figure 2B; type I in Figure 6B). The partial pairing was also sometimes observed between individual univalents (Figure 2B). Using FISH with SatCE1 DNA marker, we distinguished signal on one bivalent and one univalent (Supplementary Figure S8A). We also confirmed crossing over loci only on bivalents, while no crossing over loci was found on univalents (Figure 3B). Second population of pachytenic oocytes included diploid oocytes with 24 bivalents (n = 28) possibly represented by *C. hankugensis* chromosomes (Figure 2C; type III in Figure 6B). Such bivalents always exhibited at least one crossing over locus (Figure 3C). In addition, we detected one signal of SatCE1 DNA marker suggesting the pairing of homologous chromosomes (Supplementary Figure S8B). Finally, third population of pachytenic oocytes included haploid oocytes with approximately 25 univalents (n = 37) possibly represented by *I. longicorpa* chromosomes (Figure 2D, type II in Figure 6B). Among these oocytes, we sometimes observed incomplete pairing between 2-3 univalents. Such chromosomal spreads had 0-3 recombination loci only in paired chromosomal parts (Figure 3D). FISH with SatCE1 DNA probe revealed one signal on a univalent (Supplementary Figure S8C).

**Figure 6.**
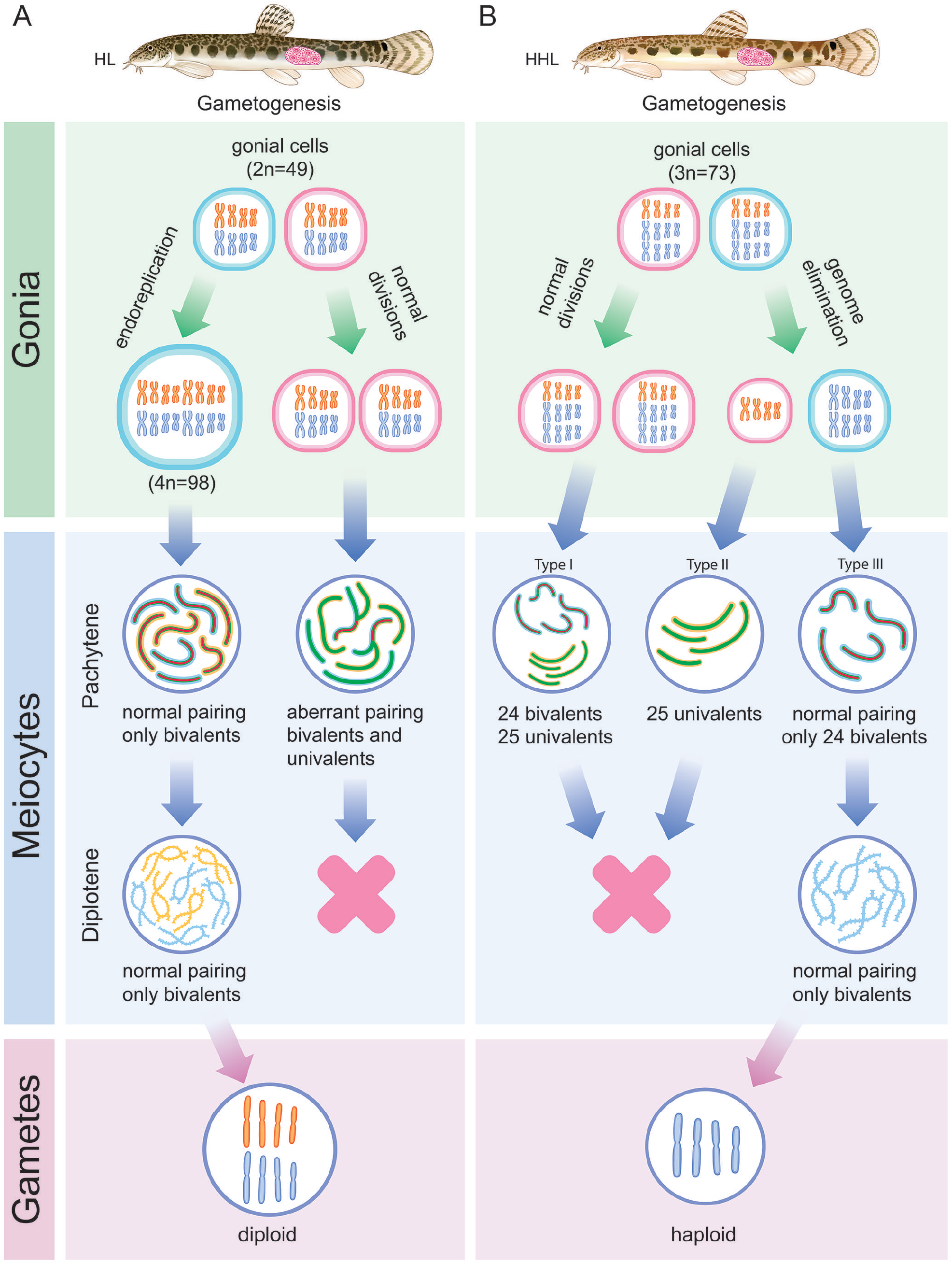
Schematic overview of gametogenesis of diploid and triploid hybrids within *C. hankugensis-I. longicorpa* complex. A. Diploid hybrids (HL) have premeiotic genome endoreplication in the minor portion of gonocytes. It allows the formation of bivalents in the pachytene stage of meiosis as each chromosome has a chromosomal copy to pair with. Afterward, such oocytes proceed to diplotene, and after meiotic completion, they form diploid gametes. B. On the contrary, in the portion of gonocytes from triploid hybrids (on example of hybrids with HHL genome composition), genome elimination of the ‘L’ genome occurs, leading to the formation of gonocytes with HH genome composition (type III) and possibly gonocytes with L genome only (type II). Most gonocytes do not have genome elimination and exist with the initial ploidy level (type I). In the pachytene stage of meiosis, type III oocytes have 24 nicely paired bivalents; type I oocytes have a mixture of 24 bivalents and 25 univalents; and type II oocytes have univalents with the partial pairing of a few chromosomes. Only type III oocytes proceed beyond pachytene into the diplotene stage of meiosis followed by the formation of reduced haploid gametes. Chromosomes of *I. longicorpa* marked in orange; chromosomes of *C. hankugensis* marked in blue; green indicates lateral elements in synaptonemal complexes; and red indicates central elements of synaptonemal complexes.

The analysis of gonadal microanatomy revealed similar distribution of gonocyte, pachytene, and diplotene oocytes in hybrids and parental species (Supplementary Figure S6D). Further, we identified ploidy in gonocytes and pachytene oocytes in intact ovary fragments using whole FISH with SatCE1 DNA markers (Figure 5A-D; Supplementary Figure S7D-F). In triploid hybrids with HHL and HLL genotypes, we discriminated gonocytes with three signals (n = 209) and with two signals (n = 26), suggesting the presence of triploid and diploid gonocyte populations correspondingly (Supplementary Table S1; Figure 5D; Supplementary Figure S7F). We assume that diploid gonocytes emerged after premeiotic genome elimination. Among pachytene oocytes, we observed cells with two signals (type I). The larger signal was localized on putative bivalents, and the small signal was detected on the dense chromatin clumps which corresponded to putative univalents (Figure 5B, C; Supplementary Figure S7E). In addition, we detected oocytes with one signal localized only on presumptive bivalent (type II). No dense chromatin clumps (presumably formed by univalent) were found in such oocytes (Figure 5B, C; Supplementary Figure S7E). We cannot distinguish between oocytes with 25 univalents and 24 bivalents based on only the FISH approach as both types of cells have single signals. However, using the morphology of chromosomes in pachytene oocytes, we suggest that oocytes with one signal include bivalents only (type III). In most diplotene oocytes (n = 352), we observed two adjacent signals similar to diplotene oocytes of parental species (Figure 5A; Supplementary Figure S7D). In 22 diplotene oocytes, we detected one signal, and in eight oocytes, we observed three signals. It may suggest methodological difficulties with merging two signals into one or that a small portion of oocytes with univalents and both with bivalents and univalents can proceed beyond pachytene. In any case, our results suggest that there is no elimination between pachytene and diplotene, and genome elimination occurs premeiotically.

### Triploid hybrid males are sterile due to aberrant pairing of chromosomes

We analysed 24 spermatocytes during pachytene from two triploid hybrid HHL males. The analysis of synaptonemal complexes using antibodies against their lateral (SYCP3) and axial (SYCP1) components revealed incomplete SCs with 6-13 properly formed bivalents (Figure 2E). In some bivalents, SYCP3 was usually localized to subtelomeric regions, while inner fragments of chromosomes lacked the SYCP3 signals (Figure 2E). The analysis of crossing over loci revealed 4-10 MLH1 loci per bivalent (Figure 3E). We conclude that hybrid males have aberrant pairing, with only a few chromosomes being able to form bivalents.

To further investigate the ability of spermatocyte I to proceed beyond metaphase I (MI) and check whether they can form spermatids and spermatozoa, we performed the analysis of gonadal microanatomy using 3D analysis. In the gonads of triploid hybrid males, we detected gonocytes, pachytene cells and large clusters of cells during metaphase I. No spermatids were observed (Supplementary Figure S6E). We also found clusters of cells with aberrant chromatin distribution and possibly apoptotic. After spindle visualization, we detected that cells during MI have misaligned bivalents or univalents. Maybe such cells cannot proceed beyond MI, causing the accumulation of such cells.

## Discussion

The present study investigated gametogenesis in diploid and triploid hybrids from *C. hankugensis-I. longicorpa* complex and demonstrated the instant switch between asexual and sexual reproduction in hybrids in dependence on their ploidy level. Specifically, inspecting the genome composition of pachytenic and diplotenic oocytes and gonial cells in natural diploid and triploid asexuals provided clear evidence that clonal and recombinant reproductive modes are dynamically altered in interspecific hybrids in relation to their ploidy level. In addition, we found reliable cytogenetic marker allowing precise recognition of genome composition and ploidy level of hybrids based on FISH mapping of earlier isolated satellite DNA marker.

### Fertility in hybrid females is rescued by specific aberrations in gametogenesis

The vast majority of meiocytes in diploid and triploid hybrid females have aberrant pairing of orthologous chromosomes leading to their arrest in pachytene (Figure 6; see below). However, both diploid and triploid hybrid females possess specific gametogenic alterations which partially rescue their fertility. In diploid hybrid females from *C. hankugensis-I. longicorpa* complex, fertility is possible due to premeiotic genome endoreplication in the portion of gonocytes (Figure 6A). Such gametogenic alteration causes the emergence of tetraploid gonocytes which are able to accomplish meiosis and form diploid clonal gametes (Figure 6A). Premeiotic genome endoreplication thus appears as very efficient mechanism to alleviate problems in orthologue pairing during meiotic prophase and simultaneously gain clonal reproduction (current data, and Dedukh et al., 2021, 2020b; Kuroda et al., 2018). Moreover, it seems to be a quite universal trait of hybrid asexual vertebrates, being observed in natural clonal lineages of loaches (Itono et al., 2006; Kuroda et al., 2018), and other fish, amphibians and reptiles (Cuellar, 1971; Dedukh et al., 2015; Dedukh et al., 2022a; Lutes et al., 2010; Macgregor and Uzzell, 1964; Majtánová et al., 2021; Stöck et al., 2002).

Triploid hybrids, however, seem unable to perform such a premeiotic genome endoreplication pathway and their fertility relies on different gametogenic alteration (Figure 6B). Specifically, investigated meiocytes of females with HHL genome composition eliminated a single-copied (*I. longicorpa*) genome, and formed bivalents between double-copied genomes (*C. hankugensis*) ensuring subsequent meiosis and formation of reduced haploid gametes (Figure 6B). In triploid hybrids with HLL genome composition, *C. hankugensis* genome was likely to be eliminated and *I. longicorpa* genome transmitted to gametes. Our observation is therefore consistent with predictions of previous crossing experiments (Kim and Lee, 1990; Ko, 2009; Lee, 1995; Saitoh et al., 2004), and may also explain the incidence of massive bi-directional introgression of mitochondrial genomes between parental species without any signs of admixis in the nucleus (Figure 1) (Kwan et al., 2019; Saitoh et al., 2004).

Such gametogenic alteration in triploid hybrids is known as meiotic or triploid hybridogenesis (Alves et al., 1998), and has been suggested to occur in several fish (Alves et al., 1998; Cimino, 1972a; Cimino, 1972b; Goddard et al., 1998; Kim and Lee, 1990; Saitoh et al., 2004) and amphibians (Dedukh et al., 2015; Graf and Polls-Pelaz, 1989; Stöck et al., 2002; Stöck et al., 2012) hybrid complexes. Nevertheless, previous data included some contrasting patterns and our observation thus provides the most comprehensive data on cellular mechanisms of genome elimination to date. Earlier studies of fish triploid hybrids of the genus *Squalius* reported pachytene cells with both univalents and bivalents, suggesting the meiotic genome elimination (Nabais et al., 2012). However, this hypothesis was based on a low number of analyzed pachytenic oocytes, probably insufficient to detect different populations of oocytes. On the other hand, in natural triploid hybrids of *Misgurnus anguillicaudatus*, premeiotic elimination of a single-copied genome was suggested based on the analysis of diplotene oocytes (Morishima et al., 2008b). However, in triploid hybrids of *Misgurnus anguillicaudatus* obtained from laboratory crosses between a sexual female and a tetraploid hybrid male, meiotic elimination of single copied genome was hypothesized (Zhang et al., 1998). Researchers suggested, that oocytes with initial ploidy level enters meiotic division I, where only bivalents can attach to the spindle, assuring their further segregation, while univalents cannot connect to the spindle and remain scattered in the ooplasm (Zhang et al., 1998). Taken together, in comparison to other publications, our data suggest that genome elimination during triploid hybridogenesis seems to have similar gametogenic mechanisms across different triploid hybrid complexes. Interestingly, it also appears that diploid and triploid hybrids may have different gametogenic alterations in dependence of their ploidy level and genome dosage, since similarl switch from gynogenesis in diploid hybrids to triploid hybridogenesis was also found in other hybrid complexes (Alves et al., 1998; Cimino, 1972b, 1972a; Goddard et al., 1998; Kim and Lee, 1990; Saitoh et al., 2004). Nevertheless, such process would also likely controlled by taxon-specific mechanisms, since in closely related *Cobitis taenia elongatoides* hybrid complex, both diploid and triploid hybrids maintain the same type of premeiotic genome endorepplication. Detailed analysis of different gametogenic stages in unrelated organisms is therefore crucial to understand the exact mechanisms and processes of alterations during the gametogenesis of hybrids.

### Gametogenetic alterations were found only in minor cell population

Interestingly, we found that both types of gametogenic alterations are particularly rare in diploid and triploid hybrids. In diploid hybrids, the premeiotic genome endoreplication occurred only in a minor portion of gonocytes while most oocytes had unduplicated genomes (Figure 6A). Since vitellogenic and early diplotene oocytes contained exclusively tetraploid genomes, we suggest that oocytes with unduplicated genomes cannot proceed beyond pachytene due to aberrant pairing, which is in good accordance with results from several other asexual hybrid vertebrates (Dedukh et al., 2021; Dedukh et al., 2022a; Newton et al., 2016; Shimizu et al., 2000). Interestingly, the ratio between duplicated and unduplicated oocytes in *C. hankugensis-I. longicorpa* hybrid females is similar to that observed in other asexual loaches (Dedukh et al., 2021; Dedukh et al., 2022a).

Surprisingly, pachytene cells of triploid hybrids also contained several populations of oocytes differing in ploidy level (Figure 6B). Thus the ability of genome elimination in triploid hybrids also seems to be restricted to only minor population of gonocytes. Moreover, we found that, at least in some gonocytes, the genome elimination occurs before meiosis, which is evident from our observation of individual diploid pachytene oocytes and diploid gonocytes (Figure 6B). In other asexuals, genome elimination may be partial or even absent during gametogenesis leading to aneuploidy in meiocytes and gametes (Chmielewska et al., 2022; Dedukh et al., 2015; Dedukh et al., 2019b). Nevertheless, we did not observe aneuploid oocytes and oocytes with univalents during diplotene. This suggests that high stringency of the checkpoint between pachytene and diplotene. Similarly, in European *Cobitis* hybrids, we earlier found that oocytes with univalents were never able to proceed beyond pachytene, possibly due to similar stringency of pachytene checkpoints (Dedukh et al., 2021; Dedukh et al., 2020b; Marta et al., 2023).

### Premeiotic genome endoreplication and genome elimination possibly occur during different ontogenetic stages

Genome endoreplication seems to be a common mechanism with probably similar underlying pathways even among unrelated lineages, however its molecular and cellular basis has not been unraveled so far. It was hypothesized that genome endoreplication might emerge in gonocytes responding to the stimulus emitted by apoptotic pachytene oocytes with unduplicated genomes (Dedukh et al., 2022a). Nevertheless, our results contrast this hypothesis at least for the studied species as HL diploid hybrids similarly to HHL and HLL triploids possess large number of pachytene oocytes with aberrant pairing that do not proceed into diplotene. However, we did not observe any sign of genome endoreplication in triploid HHL and HLL females. Thus, we incline to earlier hypothesis suggesting that aberrations in cell cycle machinery caused by hybridization may affect the cell cycle in hybrids, causing genome endoreplication (Dedukh et al., 2021). This hypothesis may also explain the absence of premeiotic genome duplication in sexual species.

Our earlier results from asexual diploid and triploid European loaches suggest that premeiotic endoreplication occurs in just one or two divisions before entering meiosis as gonocytes and pachytene oocytes with duplicated genomes are rare and do not organize in clusters (Dedukh et al., 2021). Similar patterns have been observed indiploid HL hybrid females (Figure 5E, F), possibly suggesting that premeiotic genome endoreplication generally occurs before entering meiosis in adult fishes.

By contrast, our data suggest that genome elimination in triploid hybrid loaches is presumably restricted to early stages of gametogenesis. Premeiotic genome elimination was previously observed in different asexual complexes such as diploid and triploid water frog hybrids (Chmielewska et al., 2018; Dedukh et al., 2020; Tunner, 1973; Tunner and Heppich, 1981), diploid carp gudgeon hybrids (Majtánová et al., 2021), *Poeciliopsis monacha lucida* hybrids (Cimino, 1972a) and in other animals with programmed DNA elimination (Dedukh and Krasikova, 2022). In hybrid and non-hybrid organisms, genome elimination occurs either gradually (Chmielewska et al., 2018; Dedukh et al., 2020; Gernand et al., 2005; Majtánová et al., 2021; Perondini and Ribeiro, 1997; Sanei et al., 2011) or simultaneously, including all chromosomes at once (Cimino, 1972a; Esteban et al., 1997; Prantera and Bongiorni, 2012). Simultaneous genome elimination was frequently accompanied by the formation of unipolar spindles assuring the attachment and further segregation of chromosomes from one of the parental species while chromosomes from one of the parental species usually form a clustered chromatin bulb (Cimino, 1972a; Esteban et al., 1997; Prantera and Bongiorni, 2012). The presence of pachytene oocytes with 25 univalents of *I. longicorpa* (type II) thus allows us to hypothesize that whole *I. longicorpa* genome is removed simultaneously into separate cells and fails to degrade. Additionally, the absence of aneuploid oocytes in pachyene and diplotene stages also provides indirect evidence for simultaneous removal of *I. longicorpa* genome.

Moreover, gradual chromosome elimination is frequently accompanied by micronuclei formation which were frequently found in the cytoplasm of gonocytes (Chmielewska et al., 2018; Dedukh et al., 2020; Gernand et al., 2005; Majtánová et al., 2021; Sanei et al., 2011). However, the cytoplasm of gonocytes from adult HHL hybrid females contained neither micronuclei nor chromatin bulbs of whole eliminating genome, further suggesting that premeiotic genome elimination may be restricted to early gametogenesis and presumably does not occur in adult animals.

Taken together, we suggest that premeiotic genome endoreplication most likely occurs one or few divisions before entering meiosis while premeiotic genome elimination may be restricted to early gametogenic stages and does not occur in adult hybrid females. However, detailed analysis of gonads during different ontogenetic stages is required to elucidate the mechanism of genome elimination in triploid hybrids.

### Pairing of orthologous chromosomes is aberrant and cause sterility in male hybrids

In contrast to hybrid females, triploid hybrid males do not exhibit either genome endoreplication or genome elimination. During the analysis of pachytene spermatocytes of triploid hybrid males, we found aberrant pairing with several bivalents, univalents and multivalents, which is similar to diploid and triploid male hybrids between European *Cobitis* species and between Japanese species of *Misgurnus* genus (Dedukh et al., 2020b; Kuroda et al., 2019). In contrast to hybrid females, spermatocytes of triploid hybrid HHL males can bypass the pachytene and enter meiotic metaphase I despite their aberrant chromosome pairing. Nevertheless, on gonadal tissue fragments, we observed only rare spermatid and sperm cells, which matches previous histological observations showing the presence of malformed spermatids of various sizes and a high number of apoptosis (Park et al., 2011). Interestingly, rare sperm were earlier found in triploid hybrid males, albeit with significantly decreased motility compared to parental species (Yun et al., 2021). This somewhat corresponds to our finding of clusters of spermatocytes in metaphase I with aberrant chromosome attachments to the spindle, possibly due to univalent and multivalent formation during meiosis I. Thus, we hypothesize that only rare spermatocytes can bypass metaphase I leading to the formation of aberrant spermatozoa.

The high sex specific bias in the triggering of asexuality may imply a role of genetic sex determination. Transplantation of spermatogonia from hybrid males into females of sexual species within European loaches hybrid complex restored their ability to endoreplicate their gonocyte genomes. In contrast, the reciprocal transplantation experiments of oogonia from hybrid females into males caused their sterility due to aberrant pairing and inability to undergo genome endoreplication (Tichopád et al., 2022). It suggests that initiation of endoreplication, at least in European *Cobitis*, is not directly connected to the genetic sex determination of the individual, but rather to the gonadal environment, being possible only in the ovary. However, in *Misgurnus* loaches, endoreplication occurred in hormonally sex-reverted hybrid males (Yoshikawa et al., 2007), which provides a somewhat contrasting interpretation that genome endoreplication in *Misgurnus* loaches does not depend on the sex of the individual but rather is genetically determined. It may suggest that premeiotic endoreplication may proceed differently, even in closely related organisms such as *Cobitis* and *Misgurnus*. Unfortunately, the exact types of genetic sex determination systems in these two groups of loaches are not yet known.

### Reproduction of hybrids in *C. hankugensis-I. longicorpa* complex relies on asexuality-sexuality cycles

The formation of clonal gametes serves as a prerequisite for asexual reproduction (Marta et al., 2023; Moritz et al., 1989; Savidan et al., 2001). It usually requires instant modifications of gametogenic pathways in hybrid progeny to overcome sterility caused by abnormal chromosomal pairing in meiotic prophase in unduplicated cells (Dedukh et al., 2021; Marta et al., 2023), which may be possible by genome elimination or genome endoreplication, which appear to emerge instantly upon hybridization in at least some taxa (Cole et al., 2010; Dedukh et al., 2019a; Dedukh et al., 2021; Marta et al., 2023; Shimizu et al., 2000). Nevertheless, successful establishment of asexual lineages requires additional alterations of gametogenic and fertilization processes (Schön et al., 2009; Stöck et al., 2021). In stable gynogenesis, the formation of diploid eggs is usually combined with its ability to eliminate the sperm genome after fertilization (Beukeboom and Vrijenhoek, 1998; Saat, 1991; Zhang et al., 2015). Studied diploid HL hybrids indeed have clonal gametogenesis and are able to produce diploid eggs (Figure 1, 6A), however, they do not form self-maintaining asexual lineage, as their eggs incorporate sperm’s genetic material, leading to the emergence of triploid hybrids (Figure 1) (Kim and Lee, 1990; Saitoh et al., 2004). Triploid hybrids seem also unable of clonal reproduction and produce recombinant gametes after eliminating a single-copied genome (Figure 1, 6B) (current data, Saitoh et al., 2004). Depending on which parental male fertilizes their gametes, this may either lead to the emergence of new clonal diploid hybrids or of an individual with nuclear genomic constitution of the parental species (Figure 1, 6B) (Lee, 1995; Ko, 2009). Thus, while clonal gametogenesis is a necessary step toward asexual reproduction, additional modifications are required to establish self-maintaining asexual lineages.

Although the clonal reproduction of hybrids effectively restricts gene flow between genomes of both parental species (Dedukh et al., 2020b), in cases like the *C. hankugensis-I. longicorpa* complex it can facilitate mtDNA exchange between parental species (Kwan et al., 2019; Plötner et al., 2008; Saitoh et al., 2004). Earlier study indeed reported extensive introgression of the mitochondrial genome of *C. hankugensis* into *I. longicorpa* individuals and vice versa with no evidence of nuclear introgression across the species boundary (Kwan et al., 2019). Similarly, the transfer of mitochondrial genome was observed in other species exploiting hybridogenetic reproduction (Goddard and Schultz, 1993; Plötner et al., 2008; Sousa-Santos et al., 2006), which may suggest potential advantage of such cyto-nuclear hybrids in expanding the habitats (Plötner et al., 2008).

In summary, the reproduction of *C. hankugensis-I. longicorpa* complex was likely triggered by hybridization, resulting in three types of hybrids (diploid and two types of triploid). Nevertheless, the stable maintenance of *C. hankugensis-I. longicorpa* complex relies on dynamic interactions between hybrids and sexual species and relies on specific modifications of gametogenic program which vary between hybrids with different ploidy levels (Figure 1, 6A, B).

## Material and Methods

### Samples studied and preparation of specimens for cytogenetic examination

The fish samples of *Cobitis hankugensis* and *Iksookimia longicorpa* and their diploid and triploid hybrids were separately collected from three different places along the Lam Stream in the province of Unbong-eup Namwon-si Jeollabuk-do in Korea according to previous distributional reports (Lee, 1992; Lee, 1995; Ko, 2009). The ethical review for the fish collection and experiment were done and approved by the Institutional Animal Care and Use Committee (IACUC) at Ewha Womans University (IACUC permission no. 15-104).

We analysed gametogenesis in five *I. longicorpa* (one male, three females, one juvenile) and seven *C. hankugensis* (three males, four females). In addition, we investigated gametogenesis in 14 triploid HHL hybrid individuals (two males and 15 females), three triploid HLL hybrid females and two diploid HL hybrid females from natural localities. Any treatment or injection was used before the investigation of female gametogenesis. Animals were anesthetized in MS222 followed by euthanasia according to standard procedures to minimize suffering. Kidneys and testes were used for mitotic and meiotic metaphase chromosome preparations. Ovaries and testes of each individual were separated into several pieces and used for pachytene chromosome preparation and for the whole-mount analysis. Additionally, ovaries were used for diplotene chromosome preparation. For whole-mount analysis, gonadal tissue fragments were fixed in 2% paraformaldehyde in 1× PBS for 90 min at room temperature (RT), washed in 1× PBS, and transferred to 96% methanol for long-term storage.

### Species and ploidy identification

Genomic composition and the type of ploidy of every investigated specimen were firstly evaluated using the morphological examination of external characters and size of red blood cells (Ko, 2009; Park et al., 2011; Yun et al., 2021) followed by PCR-sequencing of a species-diagnostic nuclear gene, *enc 1* (ectodermal-neural cortex I), and mitochondrial cyt *b* gene (cytochrome *b*) according to previous genetic studies involving the two species (Kwan et al., 2018; Kwan et al., 2019). In other words, the ploidy level and genomic composition of each fish examined for the cytogenetic study were determined by combining two complementary methods: size measurement of erythrocytes (Ko, 2009) and DNA sequencing of the *enc 1* gene. Consequently, this combination allowed us to diagnose them unambiguously (Supplementary file S1###). The chromatogram of DNA sequences of the *enc 1* gene could let us know whether the gene sequences are the mixture of the chromosomes of *C. hankugensis* (hereafter ‘H’ type) and *I. longicorpa* (hereafter ‘L’ type). Furthermore, the chronograms of gene sequences, *enc 1*, could discriminate diploid (HL) from triploid (HHL) hybrids on the basis of the different ratios of heterozygous peaks at their variable nucleotide sites. Finally, all the genetic diagnoses of the ploidy level among hybrids were checked by the size of the erythrocytes.

### Mitotic and meiotic metaphase chromosome preparation

Mitotic and meiotic metaphase chromosome spreads were obtained from the kidneys and testes of sexual and hybrid males without colchicine treatment according to standard procedures (MacGregor and Varley, 1983). The kidneys and testes were placed in distilled water for 30 min, followed by fixation in a 3:1 (ethanol:glacial acetic acid) solution. A fixative solution was exchanged three times. Tissues were stored at 4°C until use. The cell suspension was obtained by placing fixed tissue fragments in 70% glacial acetic acid for 1 min, during which it was intensively macerated. The suspension was dropped onto slides heated to 60°С and distributed throughout the slide surface. The excess cell suspension was removed, and interphase nuclei and metaphase chromosomes remained on the slide surface after liquid evaporation. Chromosomes in metaphase were stained with 5% Giemsa solution for 10 min at RT to confirm the number, morphology and bivalent formation.

### Pachytene chromosomes and immunofluorescent staining

Spreads of synaptonemal complexes (SC) during the pachytene stage of meiosis were prepared using protocols described by (Araya-Jaime et al., 2015) and (Moens, 2006). After manual homogenization of female gonads, 20 µl of cells suspension was dropped on SuperFrost® slides (Menzel Gläser), followed by the addition of 40 µl of 0.2 M Sucrose and 40 µl of 0.2% Triton X100 for 7 min. Further, cells were fixed for 16 minutes by adding 400 µl of 2% PFA and placed vertically to remove the liquid excess. In the case of males, after homogenization of testes, 1 µl of suspension was placed into the drop (30 µl) of hypotonic solution (1/3 of 1× PBS) preliminary dropped on SuperFrost® slides (Menzel Gläser) for 20 minutes. Afterward, cells were placed vertically in 2% paraformaldehyde (PFA) for 4 minutes. Subsequently, slides with males and females SCs were washed in 1× PBS slides for 5 minutes and stored in 1× PBS until immunofluorescent staining of synaptonemal complexes was performed.

Lateral components of SCs were detected by rabbit polyclonal antibodies (ab14206, Abcam) against SYCP3 protein while the central component of SC was detected by chicken polyclonal SYCP1 (gift from Prof. Sean Burgess; (Blokhina et al., 2019)). Using a combination of SYCP3 and SYCP1 antibodies, it is possible to distinguish bivalents from univalents, as SYCP3 is localized on both bivalents and univalents while SYCP1 is accumulated only on bivalents (Blokhina et al., 2019; Dedukh et al., 2020b). Recombination loci were detected by antibodies against the MLH1 (ab15093, Abcam) proteins. Fresh slides were incubated with 1% blocking reagent (Roche) in 1× PBS and 0.01% Tween-20 for 20 min, followed by adding primary antibody for 1h at RT. Slides were washed three times in 1× PBS at RT and incubated in a combination of secondary antibodies (Cy3-conjugated goat anti-rabbit IgG (H+L) (Molecular Probes) and Alexa-488-conjugated goat anti-mouse IgG (H+L) (Molecular Probes) diluted in 1% blocking reagent (Roche) on 1× PBS for 1h at RT. Slides were washed in 1× PBS and mounted in Vectashield/DAPI (1.5 mg/ml) (Vector, Burlingame, Calif., USA).

### Diplotene chromosomes

Diplotene chromosomal spreads (also known as “lampbrush chromosomes”) were microsurgically isolated from females of parental species as well as diploid and triploid hybrids according to an earlier published protocol (Dedukh et al., 2021; Dedukh et al., 2020b). After dissection, ovaries from unstimulated females were placed in the OR2 saline (82.5 mM NaCl, 2.5 mM KCl, 1 mM MgCl2, 1 mM CaCl2,1mM Na2HPO4, 5 mM HEPES (4-(2-hydroxyethyl)-1-piperazineethanesulfonic acid); pH 7.4). Oocyte nuclei were isolated manually using jeweler forceps (Dumont) in the isolation medium “5:1” (83 mM KCl, 17 mM NaCl, 6.5 mM Na2HPO4, 3.5 mM KH2PO4, 1mM MgCl2, 1 mM DTT (dithiothreitol); pH 7.0–7.2). Oocyte nuclei were transferred to one-fourth strength “5:1” medium (called “1:4” medium) with the addition of 0.1% PFA and 0.01% 1M MgCl2 (1:4 saline medium), in which the nucleus membrane was removed, and nucleoplasm was released into the solution. Nucleoplasm was transferred into glass chambers attached to a slide filled in a “1:4” saline medium. This method ensures that each chamber contains chromosomal spread from the individual oocyte. The slide was subsequently centrifuged for 20 min at +4°C, 4000 rpm in a centrifuge equipped with Swing Bucket Rotor for slides, fixed for 30 min in 2% PFA in 1× PBS, and post-fixed in 50% ethanol for 5 minutes and 70% ethanol overnight (at +4°C).

### Fluorescence *in situ* hybridization and whole mount fluorescence *in situ* hybridization

Probes for fluorescent in situ hybridization (FISH) procedures were selected according to earlier published data (Marta et al., 2020). We applied all earlier developed probes to satDNA repeats (satCE1-satCE7) for *C. elongatoides* (Marta et al., 2020). Probes were labelled with biotin and digoxigenin by PCR using *C. hankugensis* and *I. longicorpa* DNA isolated from muscle tissue using the Dneasy Blood &Tissue Kit (Qiagen) according to the manufacturer’s protocol.

The hybridization mixture (50% formamide, 10% dextran sulfate, 2× ЅЅС, 5 ng/μl labeled probe, and 10–50-fold excess of salmon sperm DNA) was dropped on slides covered with cover slides and carefully mounted on the edges by rubber cement. We performed common denaturation of the probe and chromosomal DNA on slides at 75°C for five minutes and incubated slides overnight at room temperature (RT) in a humid chamber. After hybridization, slides were washed three times in 0.2× SSC at + 44°C for 5 minutes each. The biotin-dUTP and digoxigenin-dUTP were detected using streptavidin-Alexa 488 (Invitrogen, San Diego, Calif., USA) and anti-digoxigenin-rhodamine (Invitrogen, San Diego, Calif., USA) correspondingly. Chromosomal DNA was counterstained with Vectashield/DAPI (1.5 mg/ml) (Vector, Burlingame, Calif., USA).

Whole-mount FISH was performed according to (Dedukh et al., 2021). After storage in 96% methanol, gonadal fragments were washed three times in 1× PBS for 15 minutes each. Afterward, tissues were impregnated wiht 50% formamide, 10% dextran sulfate, and 2×SSC for 3-4 hours at 37°C. After that, tissues were placed in a hybridization mixture of 50% formamide, 2× SSC, 10% dextran sulfate, 20 ng/µl probe, and 10 to 50-fold excess of salmon sperm DNA. Gonadal tissues were denatured at 82°C for 15 minutes and incubated for 24 hours at RT. Tissues were washed in three changes of 0.2× SSC at 44°C for 15 minutes each and blocked in 4×SSC containing 1% blocking reagent (Roche) in 4× SSC for 1 hour at RT. The biotin-dUTP and digoxigenin-dUTP were detected using streptavidin-Alexa 488 (Invitrogen, San Diego, Calif., USA) and anti-digoxigenin-rhodamine (Invitrogen, San Diego, Calif., USA) correspondingly. The tissues were stained with DAPI (1 mg/ml) (Sigma) diluted in 1× PBS at RT overnight.

### Whole-mount immunofluorescence staining

Whole-mount immunofluorescent staining was performed according to the previously published protocol (Dedukh et al., 2021). Prior to immunofluorescent staining, gonadal fragments were permeabilized in a 0.5% Triton X100 in 1× PBS for 4-5 hours at RT, followed by washing in 1× PBS at RT. After incubation in a blocking solution (1% blocking solution (Roche) dissolved in 1× PBS) for 1-2 hours, tissues were transferred into a new blocking solution with the addition of primary antibodies. We used mouse monoclonal antibodies against alfaltubulin (ab7291; Abcam). Tissues were incubated with primary antibodies overnight at RT. Anti-mouse antibodies conjugated with Alexa-488 fluorochrome (Invitrogen) were applied for 12 hours at RT. Primary and secondary antibodies were washed in 1× PBS with 0.01% Tween (ICN Biomedical Inc) for 5 minutes with shaking. Tissues were stained with DAPI (1 µg/µl) (Sigma) overnight in 1× PBS at RT.

### Wide-field, fluorescence and confocal laser scanning microscopy

Tissue fragments were placed in a drop of DABCO antifade solution containing 1 mg/ml DAPI. Confocal laser scanning microscopy was carried out using a Leica TCS SP5 microscope based on the inverted microscope Leica DMI 6000 CS (Leica Microsystems, Germany). Specimens were analysed using HC PL APO 40х objective. Diode and argon lasers were used to excite the fluorescent dye DAPI and Alexa488 fluorochrome, respectively. The images were captured and processed using LAS AF software (Leica Microsystems, Germany).

Meiotic chromosomes after FISH and IF were analysed using Provis AX70 Olympus microscopes equipped with standard fluorescence filter sets. Microphotographs of chromosomes were captured by CCD camera (DP30W Olympus) using Olympus Acquisition Software. Microphotographs were finally adjusted and arranged in Adobe Photoshop, CS6 software; Corel Draw GS2019 was used for scheme drawing.

## Acknowledgements

Authors would like to thank Antonina Maslova (Saint Petersburg State University) for the help with 3D imaging and Vladislav Vasiulin for the help with the preparation of illustrations.

## Data availability

The authors state that all data necessary for confirming the conclusions presented in the article are represented fully within the article.

